# The genome of an enigmatic sea urchin parasite *Echinomermella matsi* Jones & Hagen, 1987 resolves its place among other invertebrate parasitic nematodes

**DOI:** 10.64898/2026.01.15.699767

**Authors:** Joseph Kirangwa, Erna King, Joanna Collins, Adam Bates, Mark Blaxter, Oleksandr Holovachov

## Abstract

We present a genome of *Echinomermella matsi* (Nematoda: Plectida: Benthimermithidae), a body cavity parasite of the green sea urchin *Strongylocentrotus* spp. commonly found along the coast of Central and Northern Norway. Three assemblies were generated, one from multiple individuals using Oxford Nanopore long read data and two from two individuals using PacBio long read data. The genome of *Echinomermella matsi* is 65 Mb long consisting of 7 chromosomes, with nematode Benchmarking Using Single Copy Orthologue (BUSCO) completeness reaching 61%. The *E. matsi* chromosome complement corresponds to the proposed Rhabditida ancestral linkage groups. Phylogenetic analyses using newly generated 18S rRNA genes and a multigene dataset consisting of BUSCO protein coding genes, supported by morphological observations of juveniles, firmly place *Echinomermella* within the nematode order Plectida, alongside nematode parasitoids of marine invertebrates, *Trophomera* or *Neocamacolaimus*. As a result, the generally free-living order Plectida includes at least three independently evolved lineages of nematodes symbiotic with various groups of aquatic and terrestrial invertebrates and with unicellular organisms. This, and the fact that Plectida is the closest sister lineage to Rhabditida as a whole, and one node away from the exclusively animal parasitic Spirurina, makes this lineage a valuable model for study of evolution of animal parasitism in the aquatic environment.

**Graphical abstract:** 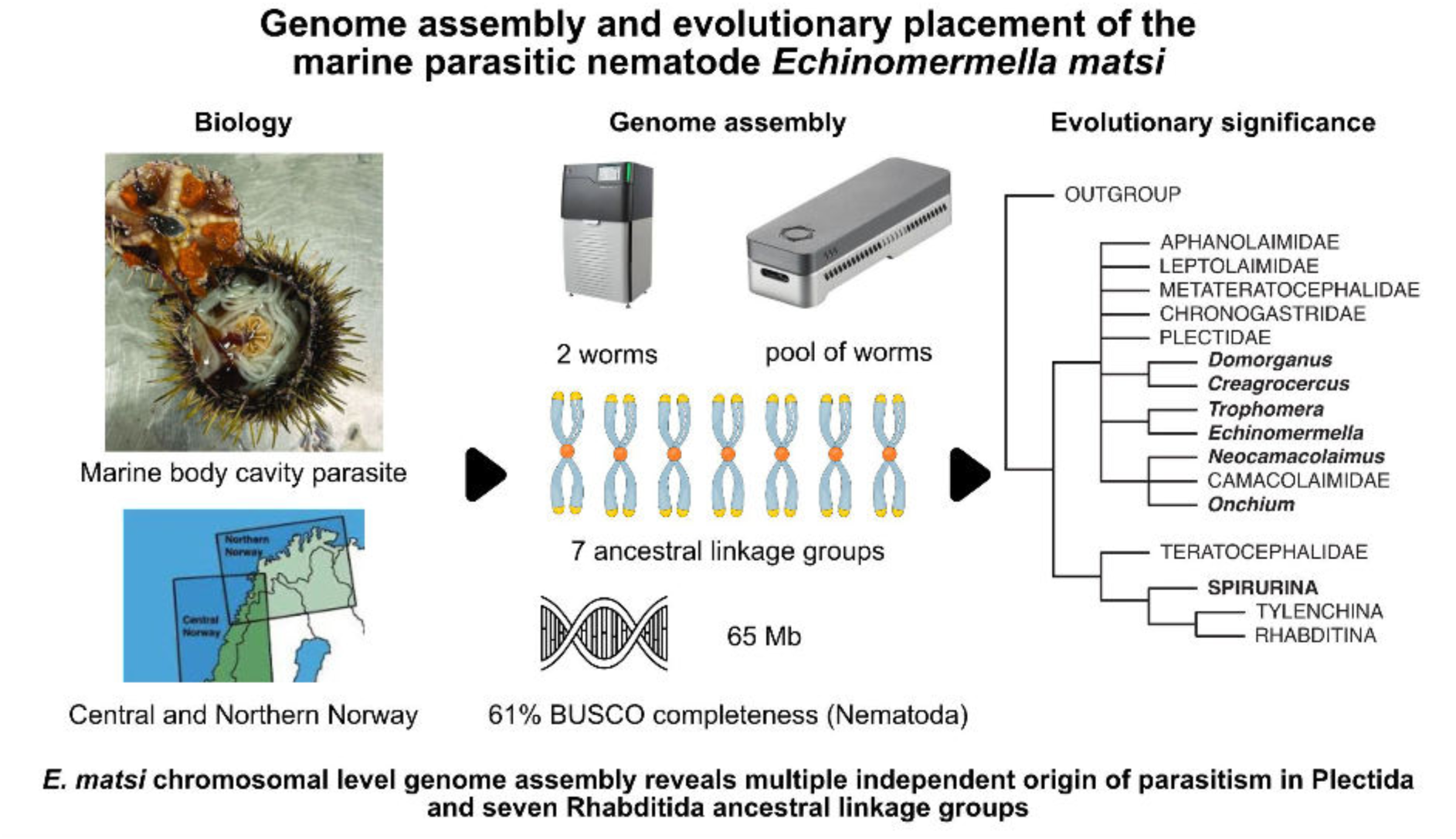

**Article summary:** The researchers generated a genome of Echinomermella matsi, a body cavity parasite of the green sea urchin, using PacBio and Oxford Nanopore long read sequencing. The genome is 65 Mb long, shows 61% of nematode BUSCO completeness, and consists of 7 chromosomes. Single and multiple gene phylogenies place Echinomermella within mostly free-living Plectida as one of the three independently evolved parasitic lineages. The authors suggest Echinomermella to be a valuable model to study evolution of animal parasitism in the aquatic environment. The genome can be used to develop biocontrol strategies of Echinomermella in mariculture of green sea urchin.

## 1. Introduction

*Echinomermella* Chitwood, 1933 are peculiar and poorly known marine nematodes. They are endoparasites of sea urchins, with only two described species. *Echinomermella grayi* (Gemmil, 1901) Chitwood, 1933, a parasite of the European edible or common sea urchin *Echinus esculentus*, was first described under the name of *Echinonema grayi* or *Ichthyonema grayi* from the Firth of Clyde in Scotland (Gemmill, 1901, Gemmill & von Linstow, 1902). The same species was subsequently recorded from the same host from several localities along the coast of Brittany (Roscoff) and the coasts of Britain (Plymouth, Scarborough, Isle of Mull and Slate Islands) and Shetland (Barel & Kramers, 1977; Comely & Ansel, 1988; Shipley, 1901), but its biology and interaction with the host remains poorly known. The second known species of *Echinomermella, E. matsi* Jones & Hagen, 1987 from the body cavity of the green sea urchin *Strongylocentrotus droebachiensis* is known much better. Originally discovered in Vestfjorden (Jones & Hagen, 1987), the species is now known to be common along the coast of Central and Northern Norway (Sivertsen, 1996), infecting over half of the population of its host in some localities (Stien et al, 1998). Naturally, such impact on a commercial species and its potential economical importance for mariculture (Hagen & Siikavuopio, 2010) has not been ignored. The morphology, distribution, ecology and host-parasite interaction of *E. matsi* have been studied in greater detail (Hagen, 1992, 1996; Poinar et al. 2011; Stien 1999; Stien & Halvorsen, 1998; Stien et al. 1996). However, the evolutionary relationships of *Echinomermella* with other animal parasitic nematodes remain unclear.

When discovered and described, these nematodes were first placed in the genus *Ichthyonema* (Gemmill & von Linstow, 1902), and later in *Philometra* (Yorke & Maplestone, 1926), other species of which are parasites of freshwater and marine fish and are currently classified in the family Philometridae (Spirurina; Clade III of Blaxter et al., 1998). Reinterpreting the data from the original description, Chitwood (1933) proposed a new genus name *Echinomermella* and provisionally placed it in Mermithoidea (equivalent to the order Mermithida, Clade I of Blaxter et al., 1998), the notion that was followed by Jones & Hagen (1987). *Echinomermella* has been associated with the parasitic “marimermithids”, a set of marine parasites now known to be polyphyletic (Tchesunov et al., 2023; Westerman et al., 2021; Zograf et al., 2024). Based on phylogenetic analysis of a single partial 18S rRNA sequence of *E. matsi* derived from a formaldehyde-preserved specimen, Poinar et al. (2011) proposed a close relationship between *Echinomermella* and marine free-living predatory *Enoplus* nematodes from the family Enoplidae (order Enoplida; Clade II of Blaxter et al., 1998). Other “marimermithid” taxa have now been placed using 18S ribosomal RNA phylogenetics within three distinct families of enoplian nematodes (Enoplia; Clade I), within Mermithida (Dorylaimia, Clade II) and within Spirurina (Rhabditida; Clade III) (Tchesunov et al., 2023). These differences led Tchesunov et al. (2023) to propose that *Echinomermella* could be “a natural model system to study genomic predispositions to parasitism”. We leverage newly sequenced genomes of *Echinomermella matsi* to redefine its phylogenetic affinities and deepen our understanding of the origin of nematode parasites of marine invertebrates.

## 2. Material and methods

### 2.1. Sampling, preservation and light microscopy

Nematodes were collected during the HHUMTL22 cruise onboard R/V Helmer Hanssen (Wernström et al., 2022). Sea urchins of unidentified species of the genus *Strongylocentrotus* were collected by divers in the Langsundet straight near Hansnes, Troms county, Norway (N 69° 57’ 40’’; E 19° 37’ 22’’) and stored in plastic containers with running seawater until dissection. Sampling did not require any state-issued permits or owner permissions. Nematodes were preserved in RNAlater, in 95% ethanol, and directly frozen at −80°C without any preservative or storage liquid. Selected specimens from ethanol were processed to absolute glycerine and mounted on permanent glass slides using paraffin as a support for the cover slip (van Bezooijen, 2006). Morphological studies were carried out using Nikon Eclipse 80i microscope equipped with differential interference contrast illumination and Sony a6400 digital camera.

### 2.2. Genome sequencing and assembly

*PacBio and Hi-C data generation and assembly:* In the Tree of Life core laboratory, tissue from the mid body of two adult females (Tree of Life identifiers neEchMats5 and neEchMats11) was homogenised individually using a PowerMasher II tissue disruptor (Denton et al., 2023). High molecular weight DNA was extracted using the Manual MagAttract protocol (Strickland et al., 2023) and purified using AMPure PB SPRI (0.45X) to eliminate shorter fragments and concentrate the DNA. The DNA concentration was assessed using a Nanodrop spectrophotometer and Qubit (dsDNA High Sensitivity Assay kit) and the fragment size distribution evaluated using the FemtoPulse system. Ultra-low input PacBio sequencing was deemed necessary for the yields obtained and DNA was therefore fragmented using the Covaris g-TUBE method (Oatley et al., 2023) and purified with AMPure PB beads (0.6X). The concentration and fragment size distribution of the sheared and purified DNA were assessed as described previously and the samples submitted to WSI Scientific Operations for sequencing on a Pacific Biosciences Revio instrument.

Hi-C libraries were prepared using the Arima-HiC v2 kit. In brief, frozen tissue (stored at –80°C) was fixed, and the DNA crosslinked using a TC buffer with 22% formaldehyde. After crosslinking, the tissue was homogenised using the Diagnocine Power Masher-II and BioMasher-II tubes and pestles. Following the kit manufacturer’s instructions, crosslinked DNA was digested using a restriction enzyme master mix. The 5’-overhangs were then filled in and labelled with biotinylated nucleotides and proximally ligated. An overnight incubation was carried out for enzymes to digest remaining proteins and for crosslinks to reverse. A clean up was performed with SPRIselect beads prior to library preparation. For Hi-C library preparation, DNA was fragmented to a size of 400 to 600 bp using a Covaris E220 sonicator. The DNA was then enriched, barcoded, and amplified using the NEBNext Ultra II DNA Library Prep Kit following manufacturers’ instructions. The Hi-C sequencing was performed using paired-end sequencing with a read length of 150 bp on an Illumina NovaSeq 6000 instrument.

The *E. matsi* neEchMats5.1 (GCA_964256745.1) and neEchMats11.1 (GCA_964248945.1) genomes were assembled following the Tree of Life Assembly process. Initial PacBio HiFi assemblies were generated with *Hifiasm* (Cheng et al., 2024), and Hi-C based scaffolding was conducted with *YaHS* (Zhou et al., 2023). The primary assemblies were analysed and manually improved using *TreeVal* (https://doi.org/10.5281/zenodo.10047653), and the chromosome-scale scaffolds confirmed by the Hi-C data were named in order of size. The mitochondrial genome was assembled using *OATK* (Zhou et al., 2025).

*Nanopore data generation and assembly.* DNA from several females was extracted using the Zymo Quick-DNA HMW MagBead Kit. A long-read sequencing library was prepared using a ligation sequencing kit SQK-LSK114 (Oxford Nanopore Technologies, Oxford, UK) following the manufacturer’s instructions. The prepared library was sequenced with R14 chemistry using MinION Mk1B and FLO-MIN114 flow cell (Oxford Nanopore Technologies, UK), and the reads were basecalled with high accuracy using *Dorado* server post-sequencing. *fastp* (Chen et al. 2018), *FastQC* (Andrews, 2024), *Meryl* (Rhie et al., 2020) and *GenomeScope* (Ranallo-Benavidez et al., 2020) were used to pre-process raw reads and assess the quality of filtered reads via the *Galaxy|Australia - Genome Lab* interface (The Galaxy Community, 2024).

The assembly, called neEchMatsONT (GCA_977018575), was generated using *Flye* (Galaxy Version 2.9.5) assembler (Kolmogorov et al. 2019) in Nanopore-HQ mode with three polishing iterations. *BlobToolKit* (Challis et al. 2020) was used to identify contigs belonging to putative contaminants and co-bionts and to evaluate the final assembly. Hi-C based scaffolding was conducted with the chromosome conformation capture data generated for the neEchMats5 specimen and using the same approaches as PacBio data (see above). Importantly, the Hi-C scaffolding and curation identified illegitimate chimaeric joins in the ONT primary assembly that had fused fragments of distinct chromosomes.

### 2.3. Phylogenetic analysis using 18S rRNA gene

Ribosomal genes were extracted from all three genome assemblies of *E. matsi* using *Barrnap* (Seemann, 2013) as implemented in *Galaxy|Australia* (The Galaxy Community, 2024), aligned and visually compared using *AliView* (Larsson, 2014). 18S rRNA from the three assemblies were identical. 18S rRNA sequences were compared with other nematode sequences using NCBI (Altschul et al. 1990) and showed no significant similarity to the existing partial 18S rRNA sequence (accession HQ668023) generated by Poinar et al (2011). To resolve the phylogenetic position of *Echinomermella*, previously published alignments (Ahmed & Holovachov, 2021) were used as templates for alignment and annotation of selected nematode sequences spanning the entire phylum, including HQ668023 and the 18S rRNA locus from the three *E. matsi* assemblies. Secondary structure annotation was manually added to all non-annotated sequences using *4SALE* (Siebel et al. 2006), and all sequences were manually aligned to maximize apparent positional homology of nucleotides. The phylogenetic tree was inferred with *IQTree* (Minh et al., 2020) using the unpartitioned alignment and built in *ModelFinder* (Kalyaanamoorthy et al., 2017) with the following command: *iqtree2 -s ./input.fas -st DNA -m MFP -b 1000 -T AUTO*.

### 2.4. Phylogenetic analysis using multigene dataset

Protein coding genes were obtained from the genome-predicted gene sets or translated transcriptomes of forty three species (including all three *E. matsi* genomes; Table 1). The protein coding ortholog genes (translated into amino acids) were extracted from each individual genome or transcriptome assembly using *BUSCO* v5.5.0 (Simão et al., 2015) and the Nematoda orthologs from OrthoDB v10 (nematoda_odb10). A custom script was used to merge identical orthologs (likely to be due to unpurged haplotypic duplication) from the genome datasets into individual orthogroup files (2,365 in total), which were subsequently aligned with *MAFFT* (Katoh et al, 2005) using the *L-INS-i* strategy. Alignments were trimmed with *ClipKIT* (Steenwyk et al., 2020) using default *smart-gap* settings to remove ambiguously aligned regions. To screen candidate core ortholog groups for paralogs and contaminants, following the strategy employed in Smythe at all. (2019) and Ahmed at al. (2022), a maximum-likelihood tree was inferred for each alignment using *IQ-TREE* (2.3.1-macOS-arm version, Minh et al., 2020) with the substitution model automatically selected by *ModelFinder* (Kalyaanamoorthy et al., 2017) for each alignment, followed by 1000 bootstrap pseudoreplicates. Next, *PhyloPyPruner* (Thalén & Kocot, 2024) was used to screen each candidate ortholog for evidence of paralogy with the following settings *--trim-lb 5 --min-support 0.75 --prune MI --mask pdist --min-taxa 25*, removing putative paralogs and contaminants, and retaining only alignments with 25 or more terminal taxa. The remaining 1,308 alignment files were used to infer maximum-likelihood gene trees with the same settings as above, using *IQ-TREE* (2.3.1-macOS-arm version, Minh et al., 2020) and substitution model automatically selected by *ModelFinder* (Kalyaanamoorthy et al., 2017), followed by 1,000 bootstrap pseudoreplicates. Coalescent-based phylogeny was inferred by combining all 1,308 gene trees using *weighted ASTRAL* (*wASTRAL*) option of *ASTER* (Zhang & Mirarab, 2022), considering clades with bootstrap support 70% and higher.

**Table 1.** Species included in the phylogenomic analysis.

| Name | BioProject/SRA accessions | Data type | BUSCO Nematoda odb10 | Source |
| --- | --- | --- | --- | --- |
| **ENOPLIDA** |  |  |  |  |
| *Enoplus brevis* | PRJEB7588 | transcriptome | C:61.2%[S:21.4%,D:39.8%],F:2.1%,M:36.7% | Koutsovoulos, 2015 |
| *Thoracostomopsis barbata* | SRR16481605 | transcriptome | C:39.1%[S:19.6%,D:19.5%],F:4.6%,M:56.3% | Tchesunov et al., 2023 |
| *Enoplolaimus lenunculus* | PRJNA953805 | genome | C:39.3%[S:36.1%,D:3.2%],F:4.5%,M:56.2% | Le et al., 2023 |
| *Pseudocella trichodes* | SRR16481604 | transcriptome | C:56.7%[S:15.9%,D:40.8%],F:3.4%,M:39.9% | Tchesunov et al., 2023 |
| *Thoracostoma sp.* | PRJNA772260 | transcriptome | C:54.3%[S:14.4%,D:39.9%],F:3.1%,M:42.6% | Tchesunov et al., 2023 |
| *Pontonema vulgare* | PRJNA504396 | transcriptome | C:62.3%[S:46.4%,D:15.9%],F:1.7%,M:36.0% | Smythe et al., 2019 |
| *Trissonchulus latispiculum* | PRJNA953805 | genome | C:35.6%[S:26.8%,D:8.8%],F:6.0%,M:58.4% | Le et al., 2023 |
| **TRIPLONCHIDA** |  |  |  |  |
| *Tobrilus sp.* | PRJNA506158 | transcriptome | C:50.1%[S:31.3%,D:18.8%],F:3.2%,M:46.7% | Smythe et al., 2019 |
| *Tripyla cf glomerans* | SRR16481603 | transcriptome | C:48.8%[S:14.5%,D:34.3%],F:4.4%,M:46.8% | Tchesunov et al., 2023 |
| **MONONCHIDA** |  |  |  |  |
| *Prionchulus punctatus* | PRJEB7585 | transcriptome | C:52.0%[S:34.0%,D:18.0%],F:6.2%,M:41.8% | Koutsovoulos 2015 |
| **MERMITHIDA** |  |  |  |  |
| *Romanomermis culicivorax* | PRJEB66727 | genome | C:39.4%[S:37.3%,D:2.1%],F:4.1%,M:56.5% | Guiglielmoni et al., 2024 |
| *Mermis nigrescens* | PRJNA802644 | genome | C:40.1%[S:35.5%,D:4.6%],F:4.4%,M:55.5% | Bhattarai et al., 2024 |
| **DORYLAIMIDA** |  |  |  |  |
| *Longidorus elongatus* | PRJEB8328 | transcriptome | C:61.3%[S:23.8%,D:37.5%],F:3.7%,M:35.0% | Danchin et al., 2017 |
| *Mesodorylaimus sp.* | PRJNA953805 | genome | C:52.4%[S:45.9%,D:6.5%],F:5.1%,M:42.5% | Le et al., 2023 |
| **DIOCTOPHYMATIDA** |  |  |  |  |
| *Soboliphyme baturini* | PRJEB516 | genome | C:35.5%[S:34.8%,D:0.7%],F:6.2%,M:58.3% | Int. Helm. Genom. Cons., 2018 |
| **TRICHINELLIDA** |  |  |  |  |
| *Trichuris muris* | PRJEB126 | genome | C:56.2%[S:55.1%,D:1.1%],F:2.5%,M:41.3% | Foth et al., 2014 |
| *Trichinella spiralis* | PRJNA257433 | genome | C:68.5%[S:38.9%,D:29.6%],F:0.7%,M:30.8% | Korhonen et al., 2016 |
| **CHROMADORIDA** |  |  |  |  |
| *Euchromadora sp.* | PRJNA506150 | transcriptome | C:39.7%[S:24.6%,D:15.1%],F:2.9%,M:57.4% | Smythe et al., 2019 |
| *Halichoanolaimus dolichurus* | PRJNSA787273 | transcriptome | C:62.3%[S:33.9%,D:28.4%],F:1.9%,M:35.8% | Ahmed et al., 2022 |
| **DESMODORIDA** |  |  |  |  |
| *Chromadoropsis vivipara* | PRJNA772260 | transcriptome | C:68.6%[S:21.4%,D:47.2%],F:2.9%,M:28.5% | Tchesunov et al., 2023 |
| *Laxus oneistus* | PRJNA609171 | transcriptome | C:70.2%[S:54.6%,D:15.6%],F:3.4%,M:26.4% | Paredes et al., 2022 |
| **ARAEOLAIMIDA** |  |  |  |  |
| *Dorylaimopsis sp.* | PRJNA767231 | transcriptome | C:64.9%[S:28.2%,D:36.7%],F:2.4%,M:32.7% | Ahmed et al., 2022 |
| *Sabatieria punctata* | PRJNA953805 | genome | C:58.5%[S:39.5%,D:19.0%],F:5.0%,M:36.5% | Le et al., 2023 |
| **MONHYSTERIDA** |  |  |  |  |
| *Linhomoeus sp.* | PRJNA953805 | genome | C:37.1%[S:28.6%,D:8.5%],F:3.8%,M:59.1% | Le et al., 2023 |
| *Paralinhomoeus sp.* | PRJNA953805 | genome | C:40.2%[S:33.0%,D:7.2%],F:4.0%,M:55.8% | Le et al., 2023 |
| *Theristus sp.* | PRJNA953805 | genome | C:42.8%[S:37.4%,D:5.4%],F:4.6%,M:52.6% | Le et al., 2023 |
| *Sphaerolaimus sp.* | PRJNA768956 | transcriptome | C:51.3%[S:15.6%,D:35.7%],F:2.5%,M:46.2% | Ahmed et al., 2022 |
| *Gammarinema scyllae* | PRJNA767233 | transcriptome | C:52.3%[S:33.5%,D:18.8%],F:1.0%,M:46.7% | Ahmed et al., 2022 |
| **PLECTIDA** |  |  |  |  |
| *Echinomermella matsi* | PRJEB79390 | genome | C:60.6%[S:60.1%,D:0.5%],F:4.7%,M:34.7% | This work (neEchMats5.1) |
| *Echinomermella matsi* | PRJEB79327 | genome | C:60.9%[S:60.4%,D:0.5%],F:4.5%,M:34.6% | This work (neEchMats11.1) |
| *Echinomermella matsi* | GCA_977018575 | genome | C:54.5%[S:54.1%,D:0.4%],F:4.0%,M:41.5% | This work (neEchMatsONT) |
| *Stephanolaimus elegans* | PRJNA768632 | transcriptome | C:31.0%[S:18.3%,D:12.7%],F:4.6%,M:64.4% | Ahmed et al., 2022 |
| *Neocamacolaimus parasiticus* | PRJNA707491 | transcriptome | C:58.5%[S:33.9%,D:24.6%],F:2.8%,M:38.7% | Ahmed et al., 2022 |
| *Anaplectus granulosus* | PRJNA506144 | transcriptome | C:38.5%[S:22.6%,D:15.9%],F:6.8%,M:54.7% | Smythe et al., 2019 |
| *Plectus sambesii* | PRJNA390260 | genome | C:75.9%[S:49.2%,D:26.7%],F:5.0%,M:19.1% | Rošić et al., 2018 |
| **RHABDITIDA** |  |  |  |  |
| *Dracunculus medinensis* | PRJEB500 | genome | C:78.3%[S:77.7%,D:0.6%],F:5.8%,M:15.9% | Int. Helm. Genom. Cons., 2018 |
| *Anguillicola crassus* | PRJNA81117 | transcriptome | C:77.0%[S:72.3%,D:4.7%],F:0.6%,M:22.4% | Heitlinger et al. 2014 |
| *Gnathostoma spinigerum* | PRJNA502990 | transcriptome | C:84.0%[S:59.5%,D:24.5%],F:3.0%,M:13.0% | Nuamtanong et al. 2019 |
| *Syphacia muris* | PRJEB524 | genome | C:82.5%[S:81.0%,D:1.5%],F:3.4%,M:14.1% | Int. Helm. Genom. Cons., 2018 |
| *Acrobelloides nanus* | PRJEB26554 | genome | C:78.8%[S:60.1%,D:18.7%],F:2.3%,M:18.9% | Schiffer et al., 2018 |
| *Bunonema sp.* | PRJNA655932 | genome | C:25.3%[S:24.9%,D:0.4%],F:2.7%,M:72.0% | Casasa et al., 2021 |
| *Poikilolaimus oxycercus* | PRJNA758215 | genome | C:54.0%[S:51.5%,D:2.5%],F:5.2%,M:40.8% | Collins et al., 2023 |
| *Rhabditoides inermis* | PRJEB71643 | genome | C:70.6%[S:63.6%,D:7.0%],F:5.3%,M:24.1% | Rödelsperger et al., 2024 |
| **NEMATOMORPHA** |  |  |  |  |
| *Gordionus montsenyensis* | PRJEB63266 | genome | C:33.4%[S:22.4%,D:11.0%],F:0.9%,M:65.7% | Eleftheriadi et al., 2024 |
| **PRIAPULIDA** |  |  |  |  |
| *Priapulius caudatus* | PRJNA20497 | genome | C:46.5%[S:37.0%,D:9.5%],F:2.1%,M:51.4% | Webster et al., 2006 |

### 2.5. Creation of figures

The three *E. matsi* genomes were mapped to the predicted rhabditid nematode ancestral linkage groups (Nigon elements) (Fig. 3) using https://pgonzale60.shinyapps.io/vis_alg/ (Gonzalez de la Rosa et al., 2021). BlobPlots were generated in *BlobToolKit* viewer v4.3.13 (Challis et al. 2020). Phylogenetic trees were visualized in *TreeViewer 2.2.0* (Bianchini, & Sánchez-Baracaldo, 2024). All bitmap images and photographs were edited using the *Affinity Photo 2* (https://affinity.serif.com/en-us/photo/) while all vector graphic illustrations and final images were created/edited using the *Affinity Designer 2* (https://affinity.serif.com/en-us/designer/) or Adobe Illustrator.

## 3. Results

### 3.1. *New data on the morphology of* Echinomermella matsi

Adult and eggs of *E. matsi* were isolated from the body cavity of *Strongylocentrotus* sp. sea urchins in northern Norway (Fig. 1A). They were identified following the original description of the species (Jones & Hagen, 1987), to which we can add some supplementary observations. Unhatched first stage juveniles possess a distinctly annulated cuticle (Fig. 1B) which is also visible in Fig. 7D in Jones & Hagen (1987). We confirm that sensory structures of the anterior end are indistinct, mostly obscured by the egg shell. The stoma of the unhatched first stage juveniles has the shape of a well defined stylet gradually extending into the lining of the pharynx, but is reduced in mature individuals (Fig. 1 C-D). The rectum in the unhatched first stage juveniles is well developed, opening to the exterior, not vestigial (Fig. 1B). The spinneret and caudal glands in the unhatched first stage juveniles are present and appear to be fully functional (Fig. 1B). Several immature females showed the remains of a rectum (Fig. 1E) appearing as a group of cells on the ventral body side, adjacent to the cuticle but without distinct lumen.

**Fig. 1.**
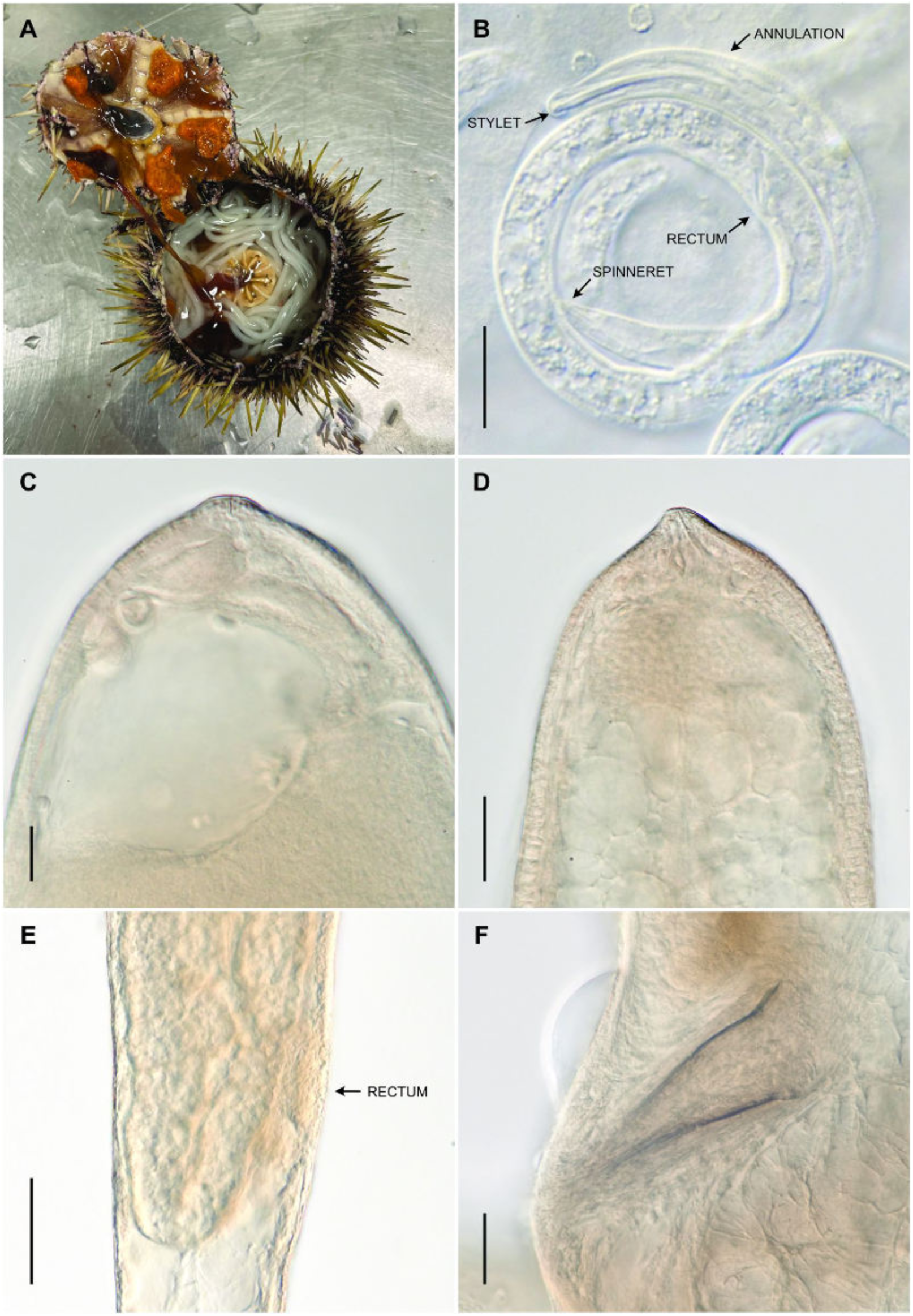
*Echinomermella matsi* Jones & Hagen, 1987. a) nematodes inside the host. b) juvenile showing a stylet, annulated cuticle, rectum and spinneret. c-d) anterior end of immature female individuals. e) posterior end of a juvenile female showing atavistic rectum. f) spicules of adult male.

### 3.2. The genome of Echinomermella matsi

The three assemblies generated have spans ranging from 52.6 to 64.6 Mb and each was scaffolded into seven chromosomes (Fig 2). The primary assembly of neEchMatsONT (Fig 2A) was more contiguous than that of neEchMats5 or neEchMats11 (Fig. 2B, C), likely because the ONT data, having a longer read N50, was able to bridge longer repeats than was the PacBio HiFi, but was reduced in span in comparison with the PacBio assemblies (Fig. 2D). This is likely because the increased error rate of ONT over that of PacBio HiFi reads leads to reduced resolution of repeat regions. The higher accuracy and longer read N50s of PacBio sequencing allows assembly and phasing of both haplotypes alongside improved resolution of multi-mapping reads over repetitive or heterozygous regions. Both PacBio assembly spans were significantly reduced upon purging of the alternate haplotypes and scaffolding to chromosomal contiguity, but retention of repeats and heterozygosity resulted in a higher final assembly span than that of the ONT assembly (Fig. 2D). We note that even in the longest PacBio assembly, neEchMats11, longer repeats such as the ribosomal RNA cistron, are still collapsed (Fig. 3A) and thus the true genome size will be larger than currently estimated.

**Fig. 2.**
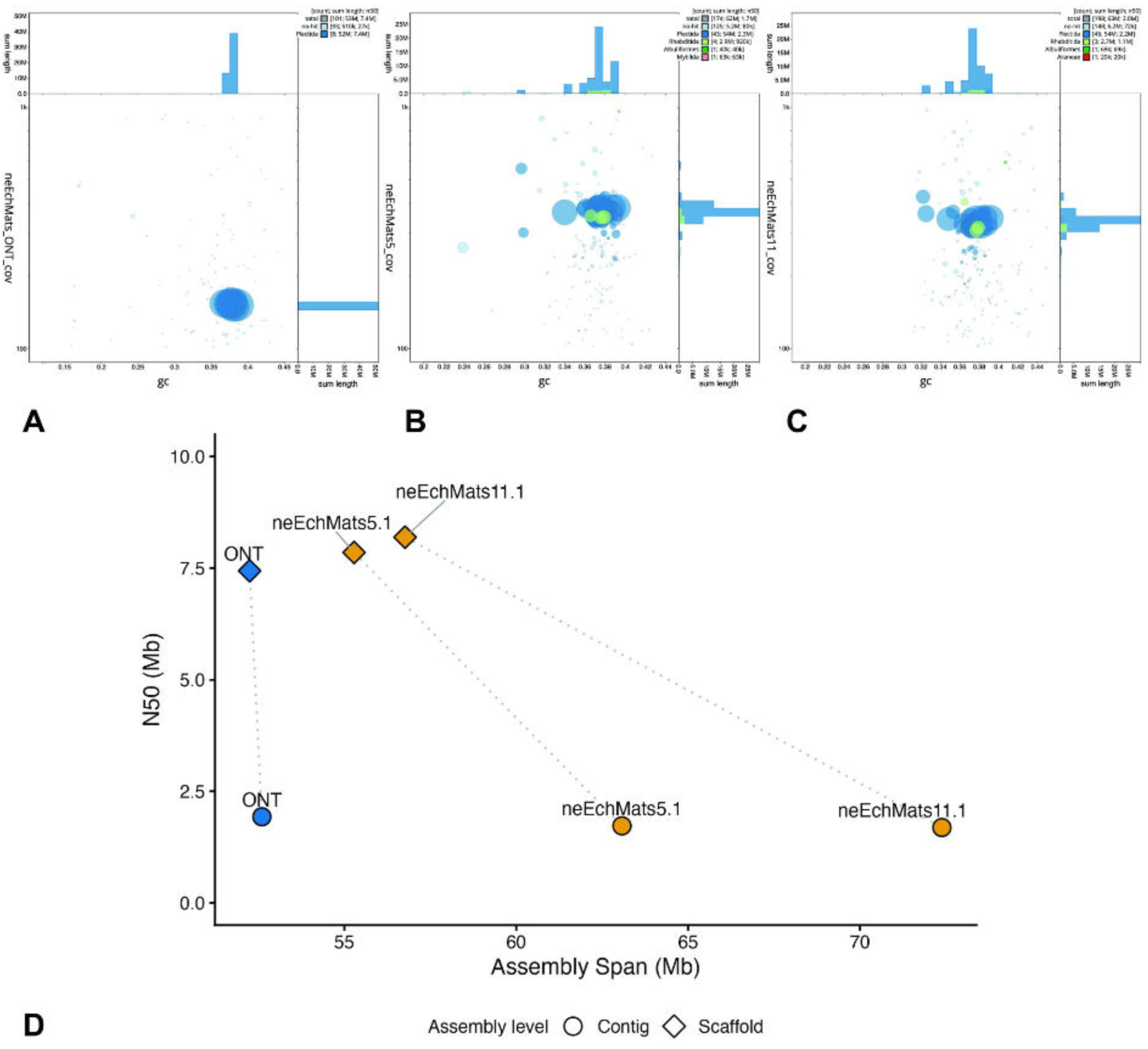
Genome assembly of *Echinomermella matsi*: a-c) BlobToolKit GC-coverage plots of primary assemblies. (a: neEchMats5.1; b: neEchMats11.1; c: neEchMatsONT). In each BlobToolKit plot, the assemblies are split into contigs, breaking scaffolds at stretches of 10 or more N residues. The x-axis shows contig GC proportion (between 0.2 and 0.45, while the y-axis shows coverage in the respective dataset. Note that coverage for neEchMatsONT is derived from mapping of the neEchMats5.1 PacBio HiFi reads. Contigs are represented by circles, with the diameter sized according to their span. Contigs are coloured by the ordinal-level taxon of the best-matching BUSCO locus on each. While most contigs are designated “Plectida”, some have aberrant best-scoring mappings to other taxa. However these mappings are not supported by more fine-scale analyses. d) Improvement in N50 and reductions in span on scaffolding with Hi-C data and removal of haplotypic duplication for the ONT (blue) and PacBio (yellow) assemblies.

Scaffolding with Hi-C resulted in resolved chromosomes for all datasets, increasing contiguity across all platforms with scaffold N50 (likely equivalent to chromosomal N50) converging around 7.8 Mb (0.4 Mb SD). The arrangement of the rhabditid nematode ancestral linkage groups (Nigon elements) (Fig. 3A) closely corresponds to the theoretical archetypal arrangement for Rhabditida (Gonzalez de la Rosa et al., 2021). Interestingly the chromosome corresponding to the predicted Nigon X included many loci previously allocated to other elements (about 50% of the loci mapped to this chromosome) while other chromosomes were more clearly assigned to one Nigon element, with <10% of loci from other Nigon element-defining groups. Chromosomes of all three individuals were highly syntenic (Fig. 3B). The increased assembly span of neEchMats11 over the other PacBio assembly neEchMats5 may reflect greater heterozygosity inherited from more divergent parents, or genuine differences in genome sizes among individuals. The higher span of neEchMats11.1 is reflected in gaps in synteny with neEchMats5.1 and neEchMatsONT on both chromosomes corresponding to Nigon element B and element N (SUPER_3 and 5 respectively), both of which are also larger in neEchMats5.1 than the ONT assembly. One potential transposition was apparent among individuals, where a Nigon E BUSCO appeared uniquely on Nigon D for neEchMats11.1 (Fig. 3 B).

**Fig. 3.**
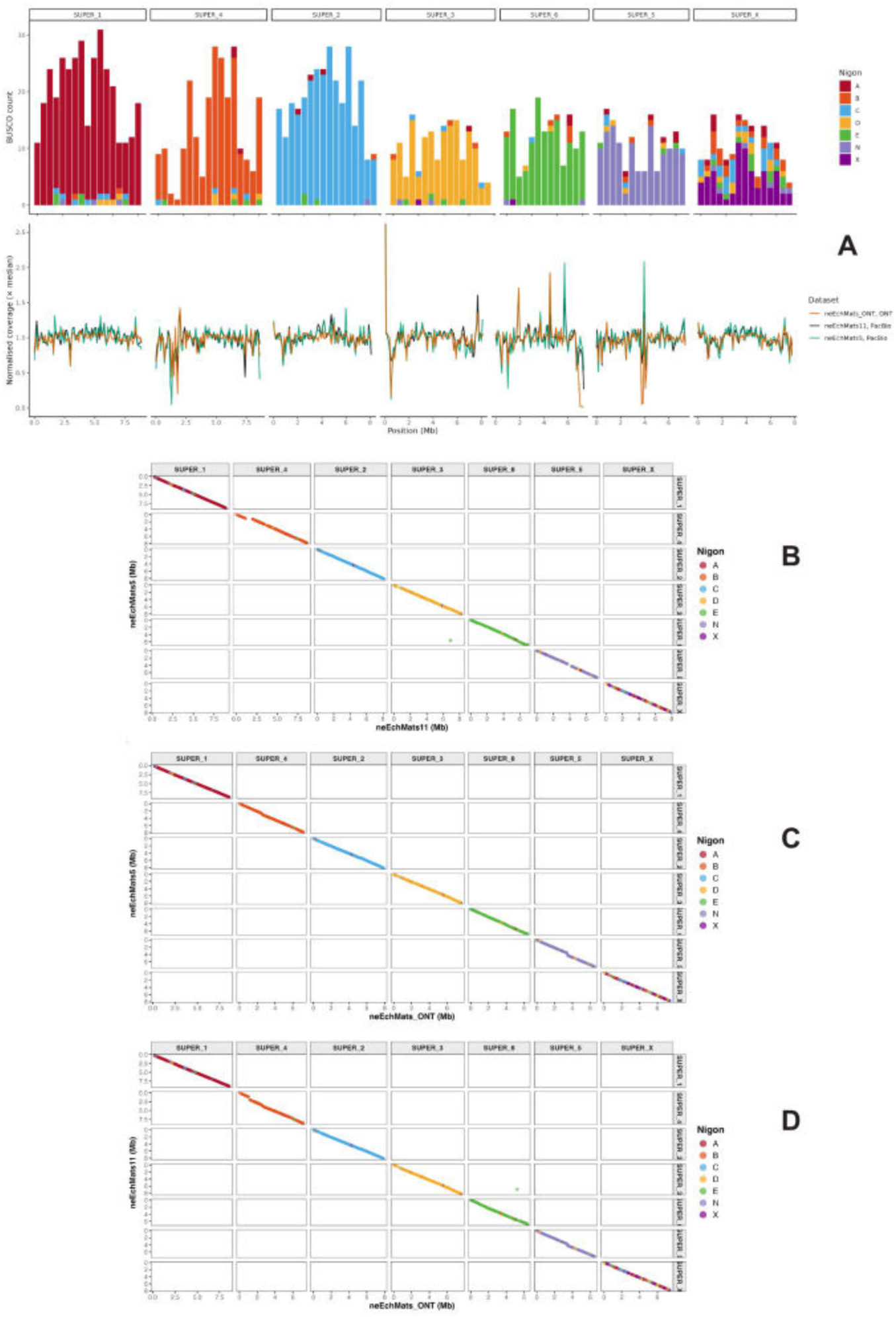
*Echinomermella matsi* retains the ancestral linkage groups of Rhabditida. a) **T**he chromosomal pseudomolecules of neEchMats11.1 correspond to intact Nigon elements. In the upper panel, for each chromosome, the count of nematoda_odb10 genes allocated to each Nigon element is plotted as a stacked histogram in 500 kb bins. The lower panel shows normalised read coverage for the data from the neEchMats5, neEchMats11 and neEchMatsONT samples. A y-axis normalised coverage limit of 2.5 was set to reduce the impact of a collapsed tandem ribosomal array at the start of SUPER_3 which had almost 5 fold normalised coverage. b) Pairwise synteny comparisons of neEchMats5.1, neEchMats11.1 and neEchMatsONT. Each chromosomal pseudomolecule is coloured by the Nigon element allocation of nematoda_odb10 BUSCO loci.

Reads for each sequenced individual were mapped against the assembly of neEchMats11.1, being the longest and highest contiguity assembly, and coverage normalised by dividing by the median to assess for differences between the sequencing platforms (Fig. 2F). Nigon X corresponds to the X sex chromosome in Rhabditid nematodes, where individuals are XX (female) and Xnull (male). We assessed read coverage of the chromosomes in our *E. matsi* read sets to identify a putative sex chromosome. The two individual nematodes sequenced (neEchMats5 and neEchMats5) were female and no difference in coverage of any chromosome was expected or observed. In the pooled nematodes sequenced to generate the neEchMatsONT assembly no drop in coverage of any chromosome was observed.

### 3.3. *Phylogenetic position of* Echinomermella matsi

BUSCO completeness assessed with the nematoda_odb10 gene set of all three assemblies ranged between 61.0 and 61.3%, with only 0.4-0.5% of the genes duplicated (Table 2). This low BUSCO completeness is not concerning, as the BUSCO nematoda_odb10 locus list is biased towards Rhabditida species, and species not included in the set used to define the list often score poorly. We used full length 18S rRNA and 1,307 nematoda_odb10 BUSCO loci in phylogenetic analyses to place *E. matsi* in the wider Nematoda phylogeny. Both phylogenetic analyses firmly place *E. matsi* within Plectida (Fig. 4), contrary to the result obtained previously using a fragment of 18S rRNA putatively from *Echinomermella* (Poinar et al. 2011, Ahmed & Holovachov 2021, Westerman et al. 2021, Tchesunov et al. 2023). This fragment (Poinar’s accession HQ668023) had no match in the *E. matsi* genome sequences and is likely to be a chance contaminant.

**Fig. 4.**
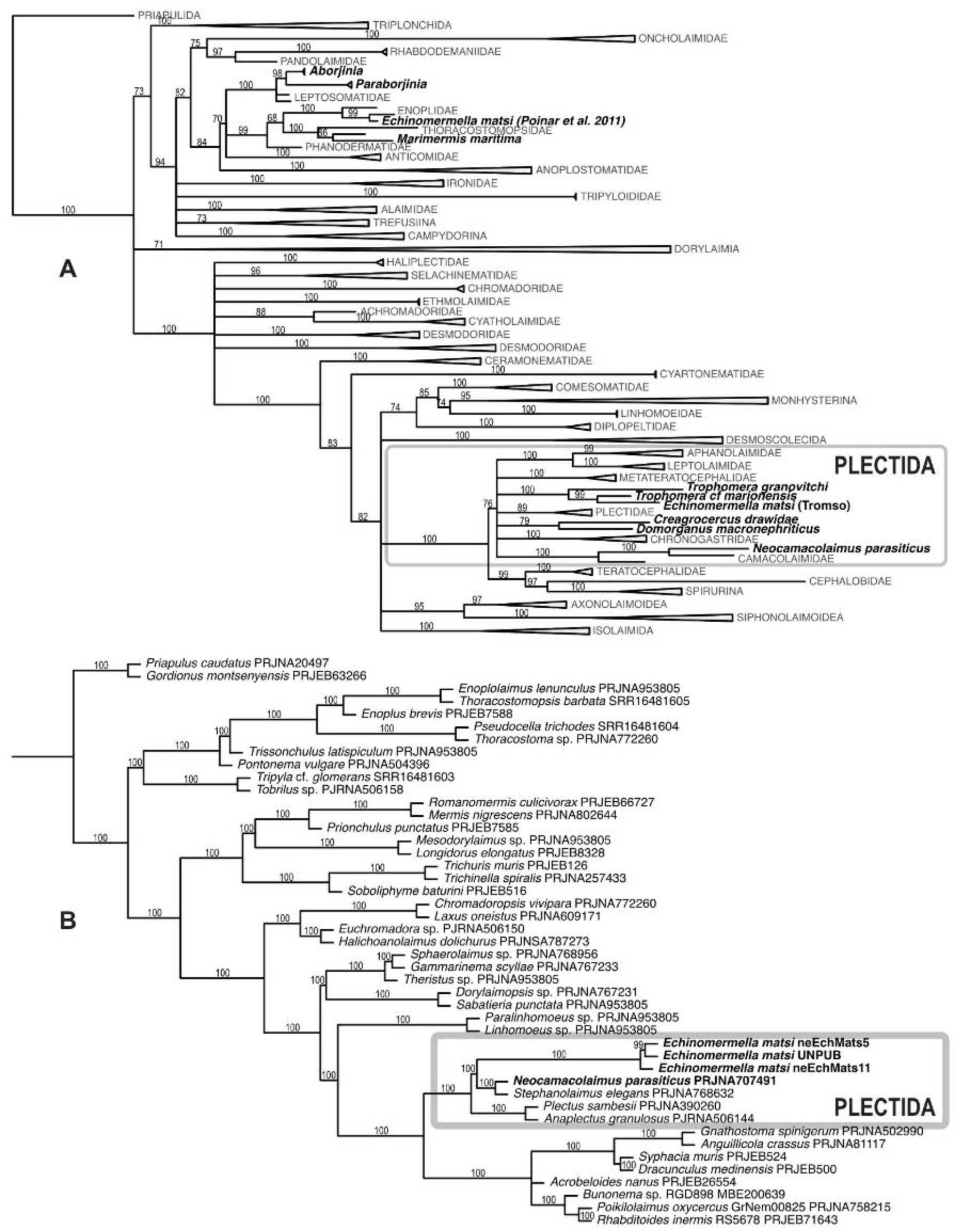
Phylogenetic relationships between *Echinomermella matsi* and other nematodes. Phylogenies were estimated using 18S rRNA (**A**) and multigene (**B**) datasets. Names in bold highlight nematodes parasites of invertebrates (mostly marine) specifically discussed in the text.

**Table 2.** Assembly statistics.

| Assembly | neEchMatsONT | neEchMats5.1 | neEchMats11.1 |
| --- | --- | --- | --- |
| Accession | GCA_977018575 | GCA_964256745.1 | GCA_964248945.1 |
| Primary decontaminated assembly length (Mb) | 52.6 | 63.1 | 72.4 |
| Contig N50 (Mb) | 1.93 | 1.72 | 1.69 |
| Contig N | 226 | 237 | 651 |
| Scaffold assembly length (Mb) | 52.2 | 55.3 | 56.8 |
| Scaffold N50 (Mb) | 7.44 | 7.85 | 8.19 |
| Scaffold N | 148 | 7 | 8 |
| BUSCO completeness * | 61.3% | 61.0% | 61.3% |
| BUSCO duplication * | 0.4% | 0.5% | 0.5% |

\* BUSCO nematoda\_odb10 gene set

The differences in taxonomic composition between the 18S rRNA locus and protein coding gene datasets result in potentially conflicting placements for *E. matsi*. The phylogenetic hypothesis based on the 18S rRNA locus includes more species, and places *Echinomermella* as an ingroup of paraphyletic *Trophomera* (Fig. 4A), a parasitoid of marine invertebrates (Tchesunov & Rosenberg, 2011), but the tree is not well resolved. The phylogeny based on 1,307 protein-coding gene trees (Fig. 4B) includes fewer species and is well resolved. *E. matsi* is placed as a sister to a clade of *Neocamacolaimus parasiticus*, another parasitoid of marine invertebrates (Holovachov § Boström, 2014) and *Stephanolaimus elegans*, a free-living marine nematode (Ditlevsen, H. 1919). *Neocamacolaimus* is placed in a separate clade from *Trophomera* in analyses based on 18S rRNA (Ahmed & Holovachov, 2020). No genomic data are yet available for *Trophomera*, but, formally the two analyses (18S rRNA and multi-gene) are not in conflict.

### 3.4. Changes to the classification of nematodes

Poinar (2011) proposed a new family Echinomermellidae and a new order Echinomermellida within Enoplia (Clade II) to accommodate this unusual parasite, in contradiction to the results of his phylogenetic analysis. Our 18S rRNA and multigene phylogenetic analyses place *Echinomermella* within the order Plectida with high support. Our 18S rRNA-based phylogeny places *Echinomermella* as a lineage within a paraphyletic *Trophomera* Rubtzov & Platonova, 1974. Based on these results we propose/confirm the following changes in the classification of the phylum Nematoda: (i) the genus *Echinomermella* Chitwood, 1933 to be placed within the family Benthimermithidae Petter, 1980; (ii) the family Echinomermellidae Poinar, 2011 to be considered a junior subjective synonym of the family Benthimermithidae Petter, 1980; (iii) the order Benthimermithida Tchesunov, 1995 to be considered a junior subjective synonym of the order Plectida Gadea, 1973; (iv) the order Echinomermellida Poinar, 2011 to be considered a junior subjective synonym of the order Plectida Gadea, 1973.

## 4. Discussion

We sequenced the genome of the enigmatic parasitic nematode *E. matsi* both to better illuminate the biology of this poorly understood marine species and to resolve conflicting concepts of the placement of *Echinomermella* within the diversity of Nematoda. Using both single-specimen PacBio HiFi sequencing and bulk-extract Oxford Nanopore sequencing, we found that these platforms were both able to generate high quality primary assemblies, but that Hi-C chromatin conformation capture data was critical in resolving the assemblies to full chromosomal level. Importantly, we find that the seven chromosomes of *E. matsi* correspond to the seven ancestral linkage groups, or Nigon elements, predicted from analysis of the rearranged genomes of rhabditid nematodes. This finding strongly supports the model proposed by Gonzalez de La Rosa et al. (2021) that the last common ancestor of Rhabditida had seven chromosomes, and suggests that this ancestral linkage group model can be extended back to the last common ancestor of Plectida and Rhabditida.

*E. matsi* is robustly placed within the generally free-living order Plectida. Plectida includes both marine and fresh-water taxa, with the bulk of species being marine detritivores. With the addition of *Echinomermella* Plectida now includes several distinct, independently evolved lineages that have parasitic relationships with invertebrates and protists. The following three monophyletic lineages can be characterized (Fig. 5):

1. A partially supported lineage uniting *Echinomermella* and *Trophomera.* Two additional monotypic genera, *Adenodelphis* Petter, 1983 and *Bathynema* Miljutin & Miljutina, 2009, are closely related to *Trophomera*. Both species of *Echinomermella* inhabit the body cavity of sea urchins as adults and both have very limited distribution compared to the areas inhabited by their hosts. The other three genera can be classified as parasitoids, as they inhabit the body cavity of their hosts as juveniles but have non-feeding, free-living adults. The precise identity of the host species is known only for some *Trophomera* species, and for *Adenodelphis*, and include nematodes, polychaetes, priapulids, molluscs, holothuroids and various crustaceans (Tchesunov & Ivanenko, 2022).
2. A monophyletic clade uniting the genera *Creagrocercus* Baylis, 1943 (family Creagrocercidae) and *Domorganus* (family Ohridiidae). All three known species of *Creagrocercus* are endoparasites of terrestrial earthworms, inhabiting the coelomic cavity of the host, but nothing else is known about their ecology or interactions with the hosts (Ivanova & Spiridonov, 2011). The biology of the genus *Domorganus* Goodey, 1946, with ten described species, is somewhat better known. Although most of its species were found in marine and freshwater sediments and soil (Holovachov, 2012), three are associated with several species of oligochaetes, inhabiting the intestine and possibly feeding on the gut microbiome (von Thun, 1967; Valovaya, 1989; Tchesunov & Sturhan, 2023).
3. The family Camacolaimidae is represented mostly by free-living species, the feeding biology of which remain enigmatic, but there are two exceptions. *Neocamacolaimus parasiticus* (Holovachov & Boström, 2014) and a similar unnamed species (Tchesunov, 2009) are parasitoids of polychaetes, where the juveniles develop in the coelomic cavity of the host, while adults (known only for *Neocamacolaimus parasiticus*, and even then only males were found) are free-living. Another, unrelated to *Neocamacolaimus,* group of species inhabit cells of foraminiferans, both testate and atestate, and include *Smithsoninema inaequale* (Hope & Tchesunov, 1999), several species of *Onchium* (Holovachov, 2015) and several unidentified and undescribed species of Camacolaimidae (Miljutin & Miljutina, 2015; Holovachov, unpublished). Observations of fixed specimens suggest that these nematodes can reproduce and develop within the foram cell, but nothing else is known about their biology.

**Fig. 5.**
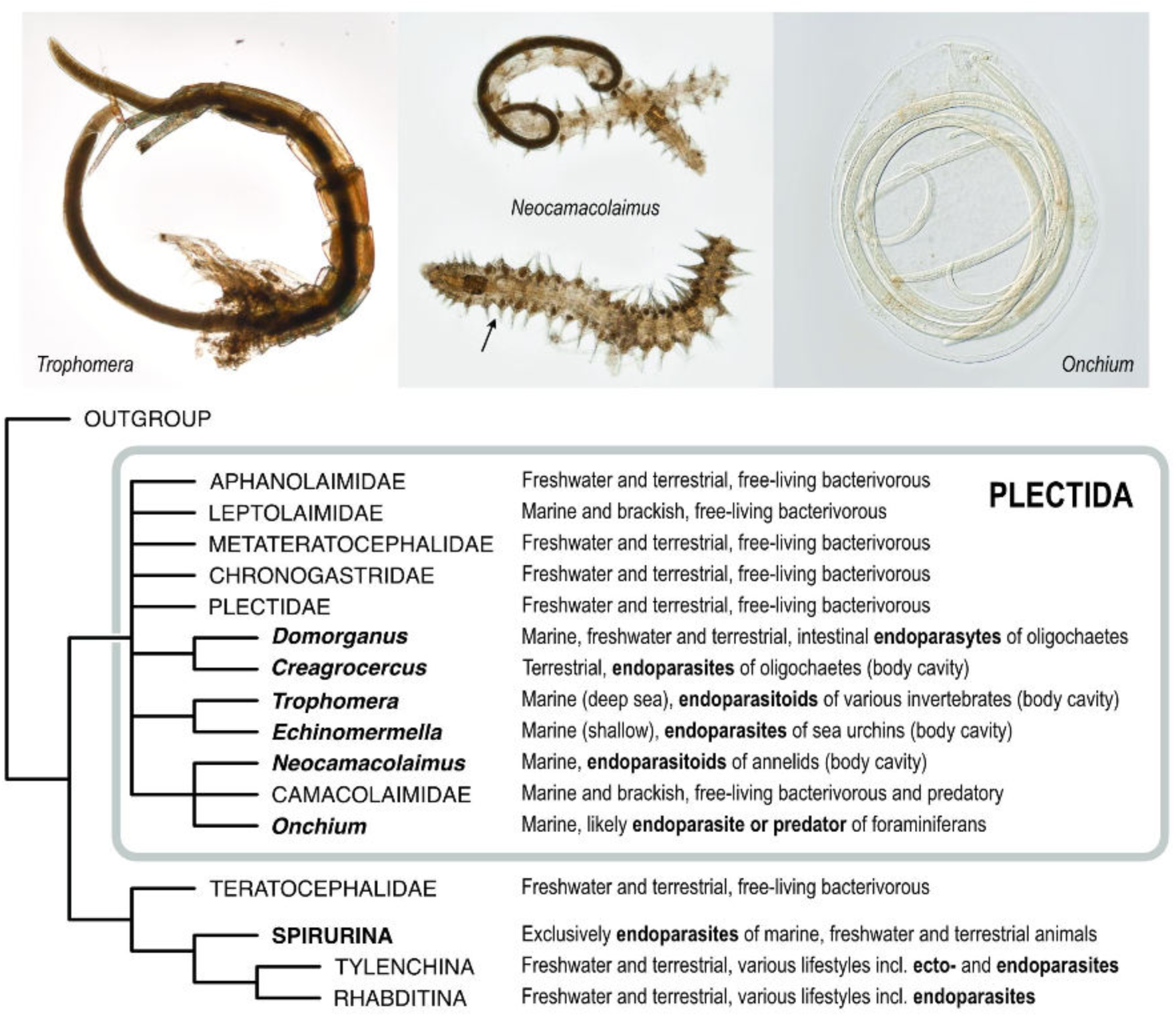
Simplified phylogeny showing known animal-parasitic lineages within Plectida. Parasites in the order Plectida and their relationships to free-living lineages and to Spirurina (Clade III) are shown, with photographic depictions of *Trophomera, Neocamacolaimus* and *Onchium* inside their respective hosts. Photographs by OH.

Parasitism has originated many times in Nematoda (Blaxter and Koutsovoulos 2014), and especially within Rhabditida. Plectida are the closest sister lineage to Rhabditida as a whole, and just one node away from Spirurina (Clade III). All species in Spirurina are parasites of animals, including important parasites of humans (the causative agents of filariasis and onchocerciasis) and farm animals (*Ascaris* large roundworms). While the phylogeny of Spirurina is not yet well resolved (Nadler et al. 2007), the earliest branching families are Gnathostomatidae and Anguillicolidae, Cucullanidae, Kathlaniidae, Quimperiidae and Seuratidae (Ahmed & Holovachov, 2021). All of these taxa use vertebrates as definitive hosts, while their intermediate hosts, when known, are crustaceans and fish. There are no known “intermediate” free-living forms with preadaptations to parasitism (Sudhaus, 2010). While two groups, Oxyurida and Rhigonematida, are parasites of terrestrial insects, they are not the earliest-branching clades in Spirurina, although their morphology and biology in general retains many features of an “average” ancestral, terrestrial rhabditid (Sudhaus, 2010).

One fundamental unanswered question is whether parasitism in Spirurina originally evolved in the marine environment or on land. Hypotheses explaining the origin of animal parasitism in terrestrial environments include preadaptations in bacterial-feeding, saprobiontic nematodes (Anderson, 1984; Sudhaus, 2010) that allow them to invade the intestine of a host, continue feeding on the gut microbiome where they withstand an anaerobic environment with higher osmotic pressure and protect themselves from digestive enzymes (in addition to other factors discussed in detail by Sudhaus, 2010). Marine invertebrates, as potential hosts, have two major differences from terrestrial counterparts (Tchesunov & Ivanenko, 2022). Most marine invertebrates do not depend on the gut microbiome to break down their food, with the possible exception of littoral oligochaetes inhabited by *Domorganus* (von Thun, 1967; Valovaya, 1989). Secondly, the body cavity of marine invertebrates is almost isotonic compared to sea water. As such, the path towards animal parasitism in marine environments must involve very different physiological and genomic adaptations compared to the same process in terrestrial habitats.

Nematodes of the order Plectida include several genera and species with well characterised genomes and transcriptomes. Some terrestrial, non-parasitic Plectids can be maintained in the laboratory and are becoming satellite models for comparative genomics (Guiglielmoni & Schiffer, 2024). Deeper study of Plectida, and especially of the independent origins of parasitism within the order, thus represents an unparalleled opportunity to shed light on the origin and early evolution of Spirurina, the most diverse and economically important group of animal parasitic nematodes.

## Data availability

Several complete and partial individuals of *Echinomermella matsi* are deposited both in ethanol and on permanent microscopic slides in the invertebrate collection of the Department of Zoology of the Swedish Museum of Natural History (accession numbers SMNH 224932–224953). Raw data used for PacBio assembly and scaffolding are available from the Sequence Read Archive (accession numbers for Revio ERX12753396, ERX12816597 and for Illumina NovaSeq X paired end ERX12760447–ERX12760450). Raw data used for ONT assembly are available from the European Nucleotide Archive (accession number ERX15144204). Assembled genomes are available from INSDC databases under BioProject PRJEB79327, accession numbers GCA_964256745.1 (neEchMats5.1) and GCA_964248945.1 (neEchMats11.1) and BioProject PRJEB97261, accession number GCA_977018575 (neEchMatsONT).

## Acknowledgements

We are grateful to Joel Burman, Sebastian Sivam Wada, Jean Baptiste Kopi and Åshild Øie Dybdal for collecting sea urchins for this project, and to Andreas Altenburger from The Arctic University Museum of Norway for mediating the process. We thank colleagues in the Scientific Operations Long Read team at the Wellcome Sanger Institute for expert support with PacBio and Hi-C data generation.

## CRediT authorship contribution statement

**Joseph Kirangwa:** Writing – original draft, Methodology, Investigation, Formal analysis. **Erna King:** Writing – original draft, Investigation, Data curation, Formal analysis. **Joanna Collins:** Investigation, Data curation. **Adam Bates:** Investigation, Data curation. **Mark Blaxter:** Investigation, Formal analysis, Project administration. **Oleksandr Holovachov:** Writing – original draft, Resources, Methodology, Investigation, Formal analysis, Data curation, Conceptualization. All authors reviewed the final manuscript.

